# A comprehensive characterization of cis-acting splicing-associated variants in human cancer

**DOI:** 10.1101/162560

**Authors:** Yuichi Shiraishi, Keisuke Kataoka, Kenichi Chiba, Ai Okada, Yasunori Kogure, Hiroko Tanaka, Seishi Ogawa, Satoru Miyano

## Abstract

Although many driver mutations are thought to promote carcinogenesis via abnormal splicing, the landscape of these splicing-associated variants (SAVs) remains unknown due to the complexity of splicing abnormalities. Here we developed a statistical framework to identify SAVs disrupting or newly creating splice site motifs and applied it to sequencing data from 8,976 samples across 31 cancer types. We constructed a catalog of 14,438 SAVs, approximately 50% of which consist of SAVs disrupting non-canonical splice sites (including the 3rd and 5th intronic bases of donor sites) or newly creating splice sites. Smoking-related signature substantially contributes to SAV generation. As many as 14.7% of samples harbor at least one SAVs in cancer-related genes, particularly in tumor suppressors. Importantly, in addition to previously reported intron retention, exon skipping or alternative splice site usage more frequently affected these genes. Our findings delineate a comprehensive portrait of SAVs, providing a basis for cancer precision medicine.

## Introduction

Comprehensive genomic characterization of multiple cancer types in large-scale genetic studies has increasingly broadened the catalog of somatic alterations that dictate cancer evolution, including single nucleotide variants (SNVs), small indels (insertions and deletions), and copy number alterations^1-3^. Moreover, it has also revealed disturbances in transcriptional regulation, such as expression changes and splicing defects that underlie cancer pathogenesis^1,2^. However, there has been only a little progress in the understanding of how somatic alterations in cancer genomes exert direct transcriptional consequences.

In cancer transcriptomes, splicing defects play important roles in many aspects of cancer development and progression^4-7^. Discovery of somatic variants affecting RNA splicing factors, such as *SF3B1* and *U2AF1*, which induce extensive alterations in RNA splicing (trans-acting regulation) in several kinds of cancers, highlights the relevance of RNA mis-splicing in cancer pathogenesis^7-9^. Another mechanism, which is the focus of this paper, is cis-acting regulation, in which somatic variants directly cause abnormal splicing of the affected gene. For example, somatic variants in canonical splice sites (highly conserved GT-AG dinucleotides at exon-intron boundaries) have long been reported to cause dysregulation of cancer-related genes^4,5^. These variants can induce different forms of abnormal splicing, such as exon skipping, intron retention, and activation of cryptic splice sites (SSs). Recent pan-cancer studies showed that SNVs causing aberrant intron retention in exon-intron boundaries are enriched in tumor suppressor genes (TSGs), especially *TP53*^10,11^. However, the complexity of splicing systems and the perplexing relationship between somatic variants and splicing alterations have limited the opportunities for systematic analyses of the extent and consequences of splicing-associated variants (SAVs), particularly those located other than at canonical sites (non-canonical sites).

To overcome these limitations, we have developed a novel algorithm, SAVNet (Splicing-Associated Variant detection by NETwork modeling), for detecting SAVs based on a list of somatic variants in a cohort and its matched RNA sequencing (RNA-seq) data using a rigorous statistical framework. Through this approach, we performed a comprehensive analysis of a large number of primary cancer samples across 31 cancer types from The Cancer Genome Atlas (TCGA), deciphering the landscape of splicing aberrations caused by cis-acting variants in human cancers.

### Overview of SAVNet framework

The overview of the proposed framework (SAVNet) is summarized in Fig. 1a. First, we collected evidence of abnormal splicing from tumor-derived RNA-seq data. Exon skipping and alternative 5’SS and 3’SS usage were extracted by capturing abnormal splicing junctions demarcated by split-aligned sequencing reads, whereas intron retention was identified by detecting sequencing reads spanning exon-intron boundaries (Fig. 1b). To obtain reliable and interpretable results, we focused exclusively on either (1) a somatic variant located at or close to an authentic exon-intron boundary (registered in the Refseq database), in which normal splicing is disrupted (SS disruption) or (2) a somatic variant located within a newly created SS inferred by an alternative SS usage event (SS creation). As demonstrated later, somatic variants and splicing alterations may have complicated relationships: a somatic variant occasionally generates different abnormal splicing events, whereas several different somatic variants sometimes cause the same splicing event. To represent these intricate relationships, we constructed a bipartite graph representing all potential associations between somatic variants and abnormal splicing events for each gene. Next, based on a probabilistic model for the number of abnormal splicing-supporting reads and the presence of a somatic variant, we deduced significant causal relationships through the evaluation of a Bayes factor incorporating a Bayesian model averaging framework^12,13^. A simulation study investigating the effect of the number of variant-splicing associations validated that the proposed framework can utilize the information from multiple associations for the sensitive identification of SAVs (Supplementary Fig. 1a, b).

**Figure 1:**
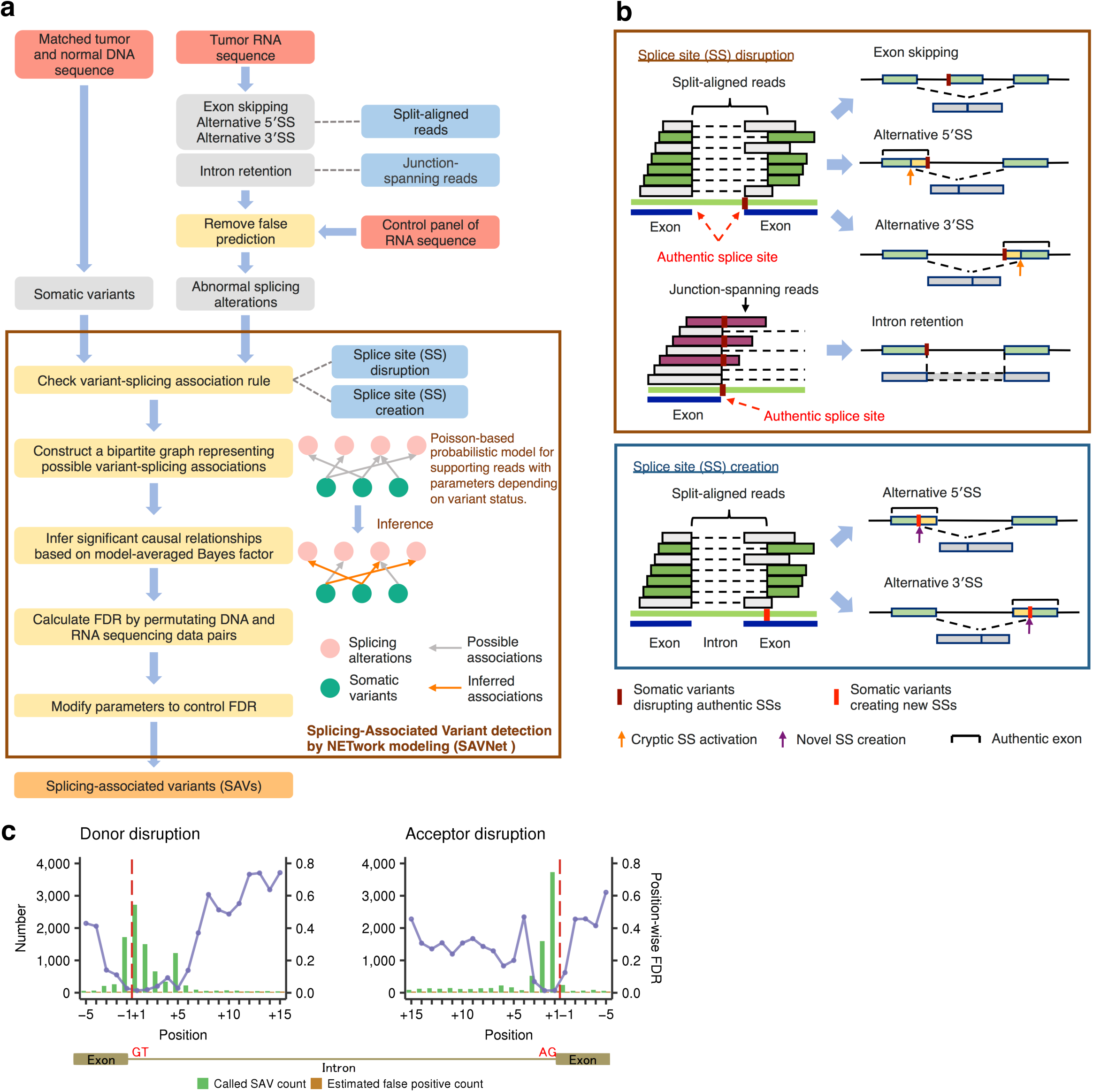
Workflow and evaluation of SAVNet. **(a)** Workflow for detecting SAVs by SAVNet from matched WES and RNA-seq data. **(b)** Schematics depicting quantification methods of exon skipping and alternative 5’SS or 3’SS usage (by split-aligned reads) and intron retention (by junction-spanning reads) and examples of somatic variants associated with abnormal splicing. SAVs within authentic SSs that disrupt normal splicing (SS disruption) and those outside authentic SSs that create alternative SSs (SS creation) were evaluated separately. **(c)** Evaluation of position-wise numbers of SAVs (green) and estimated false positives (brown) between the fifth exonic base (-5) and the 15th intronic base (+15) for splicing donor and acceptor sites. Purple points with lines show estimated position-wise FDRs. Red dashed lines represent exon-intron boundaries. See also Supplementary Fig. 1.

In the TCGA cohort, we compiled a total of 4,825,046 SNVs and 523,236 indels from 8,976 samples across 31 cancer types that underwent both whole-exome sequencing (WES) and RNA-seq using our in-house pipeline (Supplementary Tables 1 and 2 and Supplementary Methods). Initially, to determine the relevant positions within authentic SSs, we applied SAVNet to these sequencing data and assessed the accuracy of SAVNet for each position by calculating position-wise false discovery rates (FDRs) using a permutation of combinations of WES and RNA-seq data. Within authentic SSs, SS-disrupting variants at positions −3 through +6 of donor sites and −1 through +6 (except for position +4) of acceptor sites had low FDR values (below 20%), whereas much higher FDRs were observed at other positions (Fig. 1c). This observation prompted us to focus on somatic variants at these positions in the subsequent analysis. In addition, to control the overall FDR at these positions below 5%, we employed a threshold of *e*^3.0^ or greater for the Bayes factor, depending on cancer type (Supplementary Fig. 1c). To evaluate the sensitivity of SAVNet under these settings, we compared our framework with a previous study consisting of six cancer types from the TCGA^10^. In the overlapping patient population (n = 929), SAVNet detected a markedly higher number of SAVs, including two-thirds of those found in the previous study (Supplementary Fig. 1d, e). These results demonstrate the excellent detectability and satisfactory accuracy of SAVNet.

### Landscape of SAVs in human cancers

With this optimized setting, we identified 14,438 somatic variants (13,414 SNVs and 1,024 indels) responsible for 18,036 splicing alterations in the TCGA samples (Fig. 2a and Supplementary Table 3). A total of 11,153 SNVs and 875 indels disrupted splicing donor (n = 6,799) or acceptor (n = 5,229) motifs, of which 4,406 SNVs and 359 indels were not located within GT-AG canonical sites. In addition, 2,261 SNVs and 149 indels were detected to create novel splicing donor (n = 1,566) and acceptor (n = 844) sites. Thus, 7,175 (49.7%) somatic variants would not be expected to be identified by conventional methods that concentrate on SAVs involving canonical sites. Although the number of SAVs per sample was generally low (median of 1), there were quite a few samples with more instances of SAVs, particularly in cancer types with high somatic variant rates, such as lung and skin cancers (Fig. 2b).

**Figure 2:**
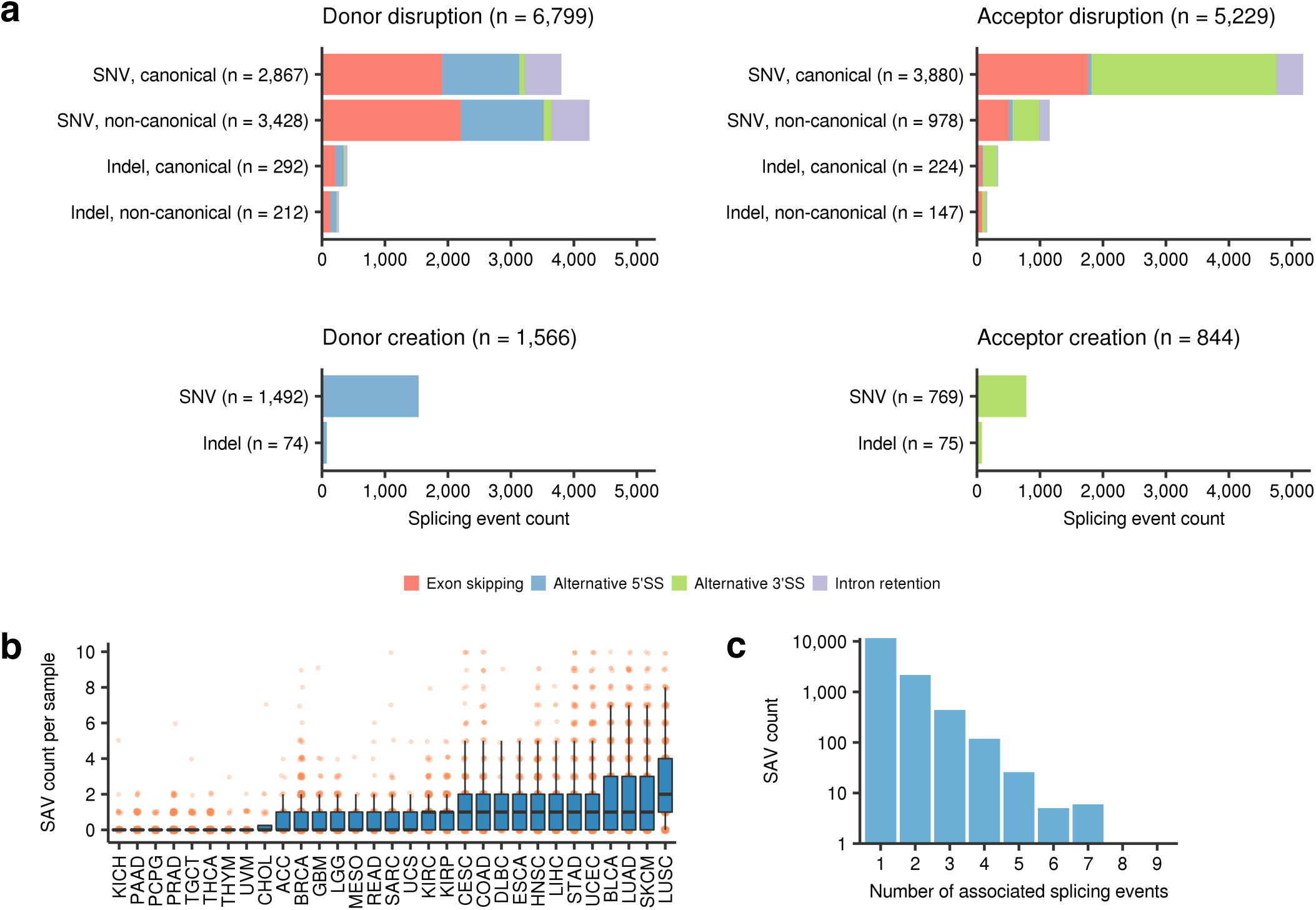
Overview of SAVs identified by SAVNet. **(a)** Number of each type of abnormal splicing events for each SAV type, stratified by (1) donor or acceptor, (2) disruption or creation, (3) SNVs or indels, and (4) canonical or non-canonical sites. Numbers in parentheses indicate the number of each type of SAV. **(b)** Number of SAVs in each sample of 31 cancer types from the TCGA project. Each point corresponds to one sample of a cancer type. **(c)** Histogram of the number of SAVs according to the number of associated abnormal splicing events. See also Supplementary Fig. 2.

Overall, these splicing alterations included exon skipping (n = 6,873), intron retention (n = 1,917), and alternative 5’SS and 3’SS usage (n = 4,522 and 4,724, respectively) (Fig. 2a). Although the vast majority of SAVs caused a single splicing alteration, 2,778 (19.2%) variants induced multiple splicing alteration events (Fig. 2c and Supplementary Fig. 2). The transcriptional consequences substantially differed according to the somatic variant pattern (donor vs. acceptor and disruption vs. creation). Exon skipping and intron retention were caused by variants disrupting both donor and acceptor sites (Fig. 2a). As expected, donor disruptions tended to generate an alternative 5’SS, whereas acceptor disruptions more frequently gave rise to an alternative 3’SS. Interestingly, exon skipping was the most frequent consequence of donor disruptions, whereas alternative 3’SS accounted for more than one-half of acceptor disruptions. Many new splice donor and acceptor sites were created by variants outside authentic SSs. Aberrant splicing events associated with variants in trans-acting splicing factors^7^ showed no overlap with those detected by SAVNet (Supplementary Tables 4 and 5).

### Positional effects of SAVs disrupting authentic SSs

To investigate the positional effects of somatic variants on splicing, we evaluated the number of SAVs disrupting authentic SSs and their ratio to overall variants according to the distance from the exon-intron boundary. This analysis revealed a substantial difference among SS positions, although the proportion of splicing outcomes was nearly consistent within donor and acceptor SSs, respectively (Fig. 3a and Supplementary Fig. 3a). As previously reported^10^, canonical GT-AG sites (at positions +1 and +2) had the highest ratios of splicing aberrations (18.3%–24.0%). In donor SSs, non-canonical sites showed a comparable total number of SAVs with canonical sites, whereas most SAVs in acceptor SSs were present at canonical sites. Together with the last exonic bases (−1) of donor sites, whose relevance was pointed out in the earlier study^10^, the fifth intronic bases (+5) also had a relatively high ratio of abnormal splicing, followed by the third intronic bases (+3). Besides GT dinucleotides, these bases are well conserved and relevant to the interaction with U1 and U5 small nuclear RNAs^14,15^. In fact, using minigene splicing assays^16^, we experimentally demonstrated that not only canonical but also non-canonical site variants cause abnormal splicing (Fig. 3b). The transcripts harboring variants at positions +5 as well as −1 showed abnormal splicing, such as exon skipping or intron retention, with comparable efficiency to canonical site variants (+1), while the wild-type transcripts were largely normally spliced.

### Mutational signatures associated with SAV generation

There were occasional discrepancies between the efficiency of somatic variants to cause abnormal splicing and the actual number of SAVs. For instance, position +2 of acceptor sites showed only a moderate number of SAVs, albeit the highest SAV ratio (Fig. 3a). These discrepancies may be attributed to the overall number of somatic variants (including those not associated with splicing alterations) and their substitution patterns at each position, which reflect both the unique base composition at SSs and the signatures of mutational processes that have been operative^17,18^. In fact, positions at −1, +1, and +5 of donor sites as well as +1 of acceptor sites, which were dominated by G bases, showed frequent G > A and G > T substitutions, suggestive of age- and smoking-related mutational processes, respectively (Fig. 3c, upper). In contrast, position +2 of donor and acceptor sites, which predominantly consist of A/T bases, showed a relatively low frequency of somatic variants. Among them, variants at canonical GT-AG sites caused splicing alterations, regardless of their base substitution pattern, whereas almost all SAVs at positions −1 and +5 of donor sites occurred at G bases, indicating almost no effect of substitutions from other bases on splicing (Fig. 3c, lower). In addition, positions having a smaller fraction of abnormal splicing were more strongly affected by the base substitution pattern. For example, G > A substitutions were the most common at position +3 of donor sites, but did not result in splicing aberrations. Moreover, despite their low frequency of overall variants, C > G substitutions (compatible with the APOBEC cytidine deaminase mutational pattern as shown below) accounted for a considerable proportion of SAVs at position +3 of acceptor sites. These findings are consistent with relatively limited conservation of splicing motifs at these positions.

**Figure 3:**
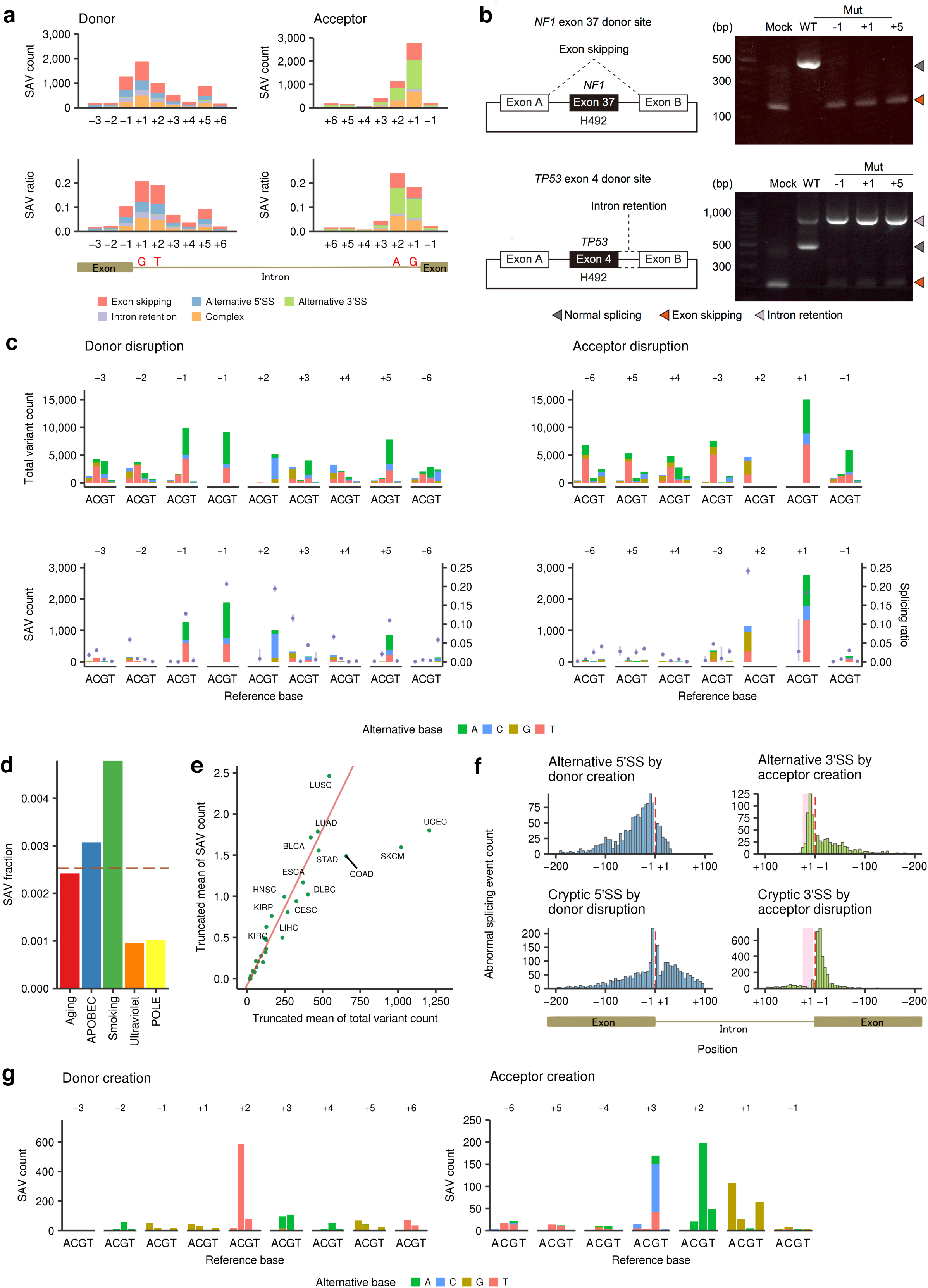
Landscape of positional differences of SAVs. **(a)** Number of SNVs disrupting splice donor and acceptor sites (SAV count, upper) and their fraction relative to total SNVs (SAV ratio, lower) at each position in the entire cohort. See also Supplementary Fig. 3a for indels. **(b)** *In vitro* splicing analyses using H492 minigene constructs (left) showing exon skipping or intron retention (right) caused by SAVs at positions −1, +1, and +5 of *NF1* exon 37 or *TP53* exon 4 donor sites, respectively. WT, wild-type; Mut, mutated. **(c)** Base substitution patterns of total somatic variants (upper) and SAVs (lower) at each exonic and intronic position of splice donor and acceptor sites. Different colors are used to display different types of alternative bases. The x-axes represent different reference bases, and the y-axes represent the numbers of variants. Fractions of SAVs relative to total somatic variants (purple points) with Bayesian confidence intervals (5% to 95% posterior quartiles) are also shown. **(d)** Fraction of estimated SAVs relative to estimated total variants attributed to each mutational signature. Red dashed line represents the overall fraction of SAVs relative to total variants. See also Supplementary Fig. 3b, c. **(e)** Scatter plot showing the relationship between SAV and total variant counts in 31 cancer types. A linear regression line (red) is fitted to the data points for each cancer, excluding those for COAD, SKCM, and UCEC. The truncated mean is used to exclude the samples with extremely large numbers of somatic variants. **(f)** Histogram showing the distribution of newly created alternative SSs (upper) and cryptic SSs caused by SS disruption (lower). Red dashed lines and pink shading represent exon-intron boundaries and polypyrimidine tract regions (positions +5 through +25), respectively. **(g)** Base substitution patterns of SAVs creating alternative SSs according to the distance from the newly created exon-intron boundaries. Colors and axes are the same as in **(c)**. See also Supplementary Fig. 3d-g.

To evaluate the underlying mutational process for each SAV, we analyzed mutational signatures found in the current sample set using pmsignature algorithm^17^. Among the five major mutational signatures (processes generating a large number of somatic variants) (Supplementary Fig. 3b), smoking signature showed the largest contribution to SAV generation, followed by APOBEC and aging signatures. Signatures related to ultraviolet exposure and altered activity of the error-prone polymerase POLE had less impact (Fig. 3d and Supplementary Fig. 3c). These differences can partly be explained by the predominance of G bases at highly affected positions (−1, +1, and +5 of donor sites and +1 of acceptor sites) and the transcriptional strand bias of several mutational signatures, i.e., the smoking signature preferentially affects G bases, whereas the ultraviolet signature frequently alters C bases. Reflecting these differences among mutational processes, lung squamous cell carcinomas (LUSC) had more SAVs than expected from the overall somatic variant rate, whereas cancers frequently affected by POLE alterations, such as uterine corpus endometrioid carcinomas (UCEC) and colon adenocarcinomas (COAD), as well as ultraviolet-associated skin cutaneous melanomas (SKCM) showed a relatively lower number of SAVs (Fig. 3e and Supplementary Fig. 3c).

### Characteristics of SAVs creating alternative SSs

Our analysis also revealed the positional distribution of SAVs creating alternative donor and acceptor sites. Newly created donor sites were widely distributed in both exons and introns, whereas abnormal acceptor sites were created predominantly within the polypyrimidine tract (Fig. 3f, upper), likely reflecting the involvement of additional conserved elements in introns, such as branchpoint sequences and polypyrimidine tracts^14,15^. Apparently similar distributions were also seen for cryptic SSs activated by variants disrupting the authentic SSs (Fig. 3f, lower). However, unlike newly created acceptor sites, a biased localization of cryptic acceptor sites toward exons was observed, which can be plausibly explained by a depletion of AG dinucleotides in the polypyrimidine tract.

We also evaluated the substitution pattern of somatic variants creating new splicing sites based on their relative position within the newly created SSs. Most newly created donor sites resulted from GT canonical site generation through C > T substitutions at position +2, whereas variants associated with acceptor creation tended to form a new YAG (Y, pyrimidine) motif at positions +3 through +1 (Fig. 3g and Supplementary Fig. 3d). These results suggest that, showing a strong bias toward particular base substitutions, these SAVs generate additional consensus donor or acceptor motifs that are more efficient for splicing than those in authentic SSs, as implicated by stronger splicing strength (assessed by MaxEnt^19^ or H-bond^20^ scores) (Supplementary Fig. 3e-g).

### Features of genomic sequences associated with SAVs

Splicing outcomes mediated by SAVs appear to be context-dependent: Somatic variants within authentic SSs can cause different forms of splicing aberration, while the same substitutions at the same relative position frequently do not alter splicing. To elucidate the factors determining the potential of somatic variants within authentic SSs to alter splicing, we compared the genomic features of SSs between normal (those not identified as SAVs) and abnormal splicing groups (SAVs) (Supplementary Fig. 4a). Generally, SS-disrupting SAVs attenuated the splicing strength more than variants that induced no abnormal splicing, regardless of the substituted position and consequent splicing alteration type (Fig. 4a and Supplementary Fig. 4b-e). A sequence motif analysis revealed a distinctive feature of SSs disrupted by SAVs, especially those at positions other than canonical GT-AG sites. As for variants occurring at the penultimate (−2) and last (−1) exon bases in the donor SSs, splicing motifs with abnormal splicing showed more conserved exonic bases but less conserved intronic bases when compared with normal splicing motifs, except for the universally conserved canonical GT dinucleotides (Fig. 4b and Supplementary Fig. 4f, g). This difference was opposite for SAVs at intronic bases, in which consensus sequences were more conserved in introns, especially at positions +4 through +6, but not in exons. These findings are compatible with the proposed mutually repressive relationship between the exonic and intronic regions of donor sites^21,22^. Analysis of the disrupted acceptor sites revealed that thymine (T) at position +4 was overrepresented in samples with SAVs at position +3, which may be due to the frequent C > G substitutions at TpC dinucleotides attributed to APOBEC activity^17,18^ (Supplementary Fig. 4f).

**Figure 4:**
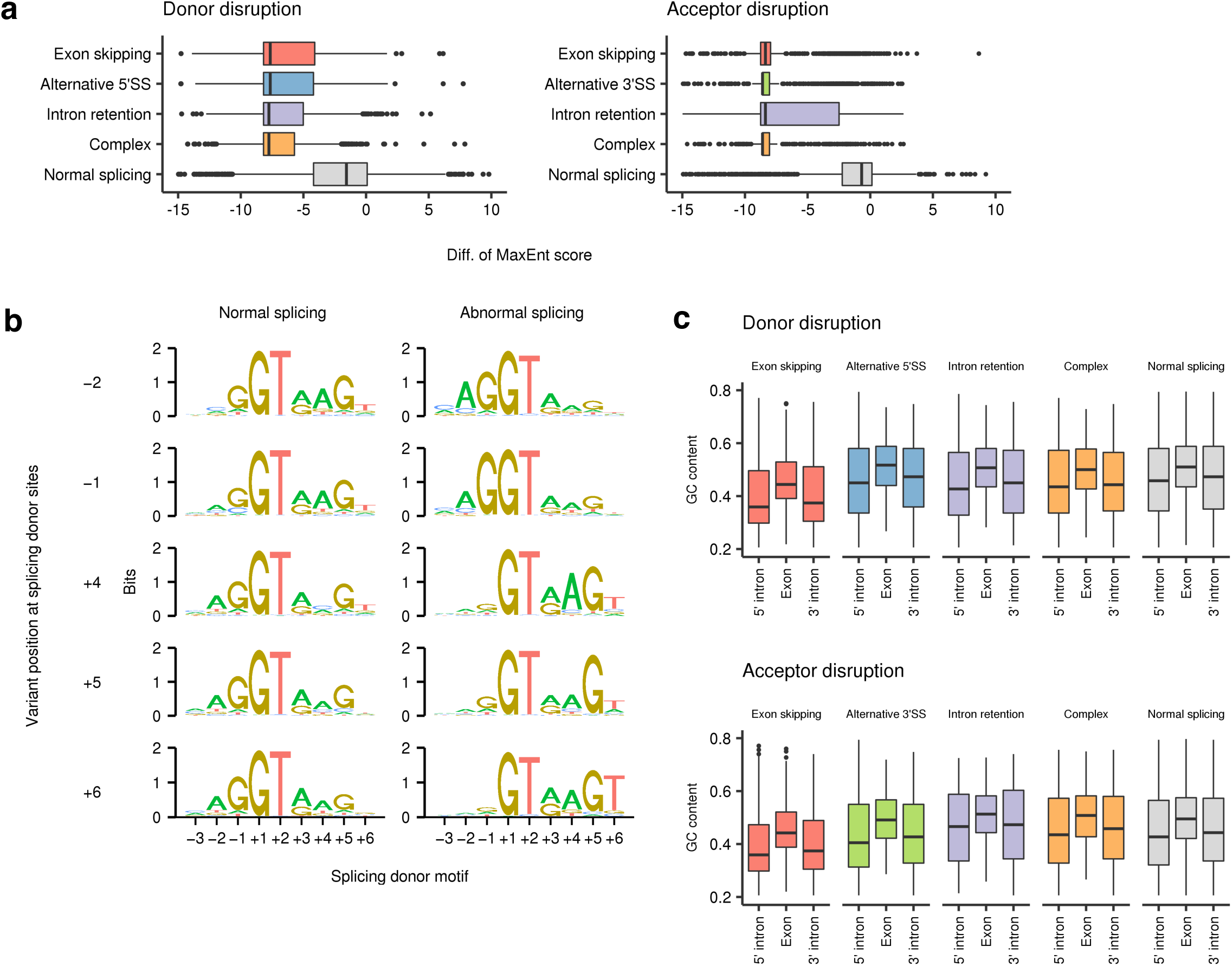
Characteristics of SS-disrupting variants generating distinct splicing alterations. **(a)** Change in splicing strength (based on MaxEnt scores) triggered by somatic variants at authentic splicing donor (left) and acceptor (right) sites according to splicing outcomes. “Complex” represents samples showing more than one splicing alteration, and “Normal splicing” represents samples lacking the relevant splicing alterations despite the presence of somatic variants in genes with detectable expression (fragments per kilobase of exon per million fragments mapped (FPKM) ≥ 10). See also Supplementary Fig. 4b-e. **(b)** Sequence motifs of splicing donor sites at which somatic SNVs lead to normal (left) or abnormal splicing (right; identified by SAVNet) according to the variant position. See also Supplementary Fig. 4f, g. **(c)** GC contents of exons affected by SAVs, and their franking 5’ and 3’ introns were compared among the five splicing groups. See also Supplementary Fig. 5.

Consistent with the previous report^10^, inspection of the exon-intron architecture revealed that exon skipping was characterized by a lower GC content in both exons and flanking introns, shorter exon and longer intron length, and stronger splicing strength (Fig. 4c and Supplementary Fig. 5a-f). These features are characteristic of SSs governed by the exon definition mechanism, in which exons are initially recognized by splicing factors^23,24^. In contrast, intron retention and alternative SS usage were associated with longer exon length, suggesting that these SSs are regulated in common by the intron definition mechanism.

### Enrichment of SAVs in TSGs

To evaluate the role of SAVs during cancer development, we investigated which genes are frequently altered by SAVs. Strikingly, the majority of frequently affected genes (present in ≥ 10 samples across the entire cohort) were well-established TSGs (Fig. 5a, b and Supplementary Fig. 6a, b). In consistent with this study^10^, in which intron retention was argued to be a major mechanism of SAV-induced TSG inactivation, SAVs that caused intron retention showed the strongest enrichment of TSGs, regardless of the cancer gene sets^1,25,26^. However, SAVs associated with exon skipping and alternative SS usage also had a greater proportion of TSGs, even when compared with nonsense variants, accounting for 88% of SAVs affecting TSGs (Fig. 5c). These findings suggest that, together with intron retention, exon skipping and alternative SS usage play crucial roles in TSG inactivation. In contrast, oncogenes were less frequently affected by SAVs, comparable to missense variants. In total, 1,684 SAVs in candidate cancer-related genes^25^ were identified in 14.7% of the TCGA samples (1,315 of 8,976). Particularly, as many as 914 SAVs found in 9.3% of samples targeted well-known TSGs^1^, of which 341 were not located at canonical sites. Moreover, SAVs accounted for 9.5% of loss-of-function variants in these genes. Therefore, SAVs represent an important but previously underestimated mechanism for TSG inactivation, irrespective of splicing outcome.

**Figure 5:**
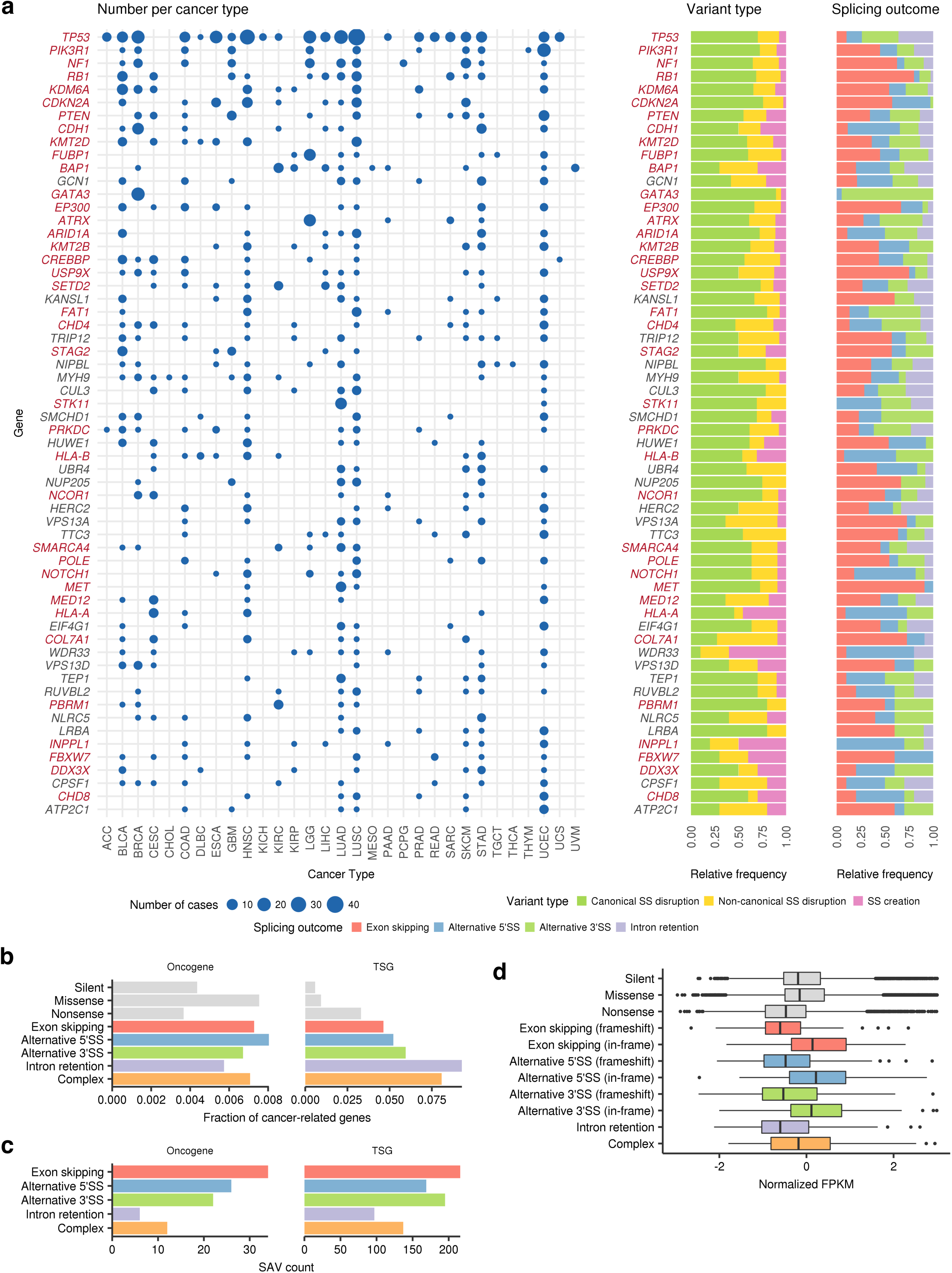
Entire spectrum of SAVs across cancer types. **(a)** Left, landscape of SAVs in frequently altered genes (total number ≥ 10) across cancer types. The point size indicates the number of affected samples. Genes are sorted by the total number of SAVs in all cancer types, and known cancer-related genes^25^ are shown in red. Right, relative frequencies of variant types and splicing outcomes of SAVs. For SAVs causing multiple splicing alterations, splicing outcomes with the largest number of supporting reads are selected. **(b)** The fractions of SAVs affecting oncogenes or TSGs (based on ref. 1) relative to total SAVs according to splicing outcomes were compared with other types of somatic variants (silent, missense, and nonsense). See also Supplementary Fig. 6. **(c)** The number of SAVs affecting oncogenes or TSGs (based on ref. 1). **(d)** Box plots showing changes in normalized (z-scored) mRNA expression (FPKM) for each splicing outcome, as compared to other types of somatic variants.

Like nonsense variants, splicing alterations are thought to trigger nonsense-mediated decay (NMD), a surveillance mechanism that selectively degrades abnormal transcripts containing a premature termination codon^6,10^. To clarify the effects of SAVs on gene expression through NMD in TSGs, we investigated the whole transcript level across different types of abnormal splicing. In line with the previous report^10^, transcripts with intron retention showed a substantially lower expression level than normal transcripts, which was comparable to those with nonsense variants (Fig. 5d). The expression of transcripts with exon skipping or alternative SS usage was also reduced when their splicing alterations caused frameshift changes.

### Genes frequently altered by SAVs

Among genes frequently targeted by SAVs, *TP53* was the most frequently altered gene, affecting 233 samples in 22 cancer types (Fig. 5a). Although the last bases of exons 4, 6, and 9 were reported to be frequently mutated^10,11^, we identified a number of recurrent variants at splice donor and acceptor sites of introns 3 through 9, with prominent base-level and/or SS-level hotspots at donor and acceptor sites of intron 4 (Supplementary Tables 6 and 7). Approximately one-half of recurrent SAVs simultaneously produced different types of abnormal splicing, while identical abnormal splicing events were generated by different SAVs, such as retention of introns 7, 8, and 9 caused by donor and acceptor SAVs of each intron (Fig. 6, upper left). Most of these SAVs induced frameshift splicing alterations, likely leading to mRNA degradation through NMD, whereas other SAVs generated in-frame exon skipping or alternative SS usage, such as exon 5 acceptor variants activating cryptic 3’SS, followed by 15-amino acid deletion. *PIK3R1*, encoding the p85 regulatory subunit of phosphatidylinositol 3-kinase, ranked second (39 samples), approximately one-half of which were found in UCEC (Fig. 5a). The majority of these SAVs caused in-frame exon skipping (mainly involving exon 4) resulting in a deletion of the iSH2 domain, which is also affected by the small deletions typically observed in this gene^27^ (Fig. 6a, upper right).

**Figure 6:**
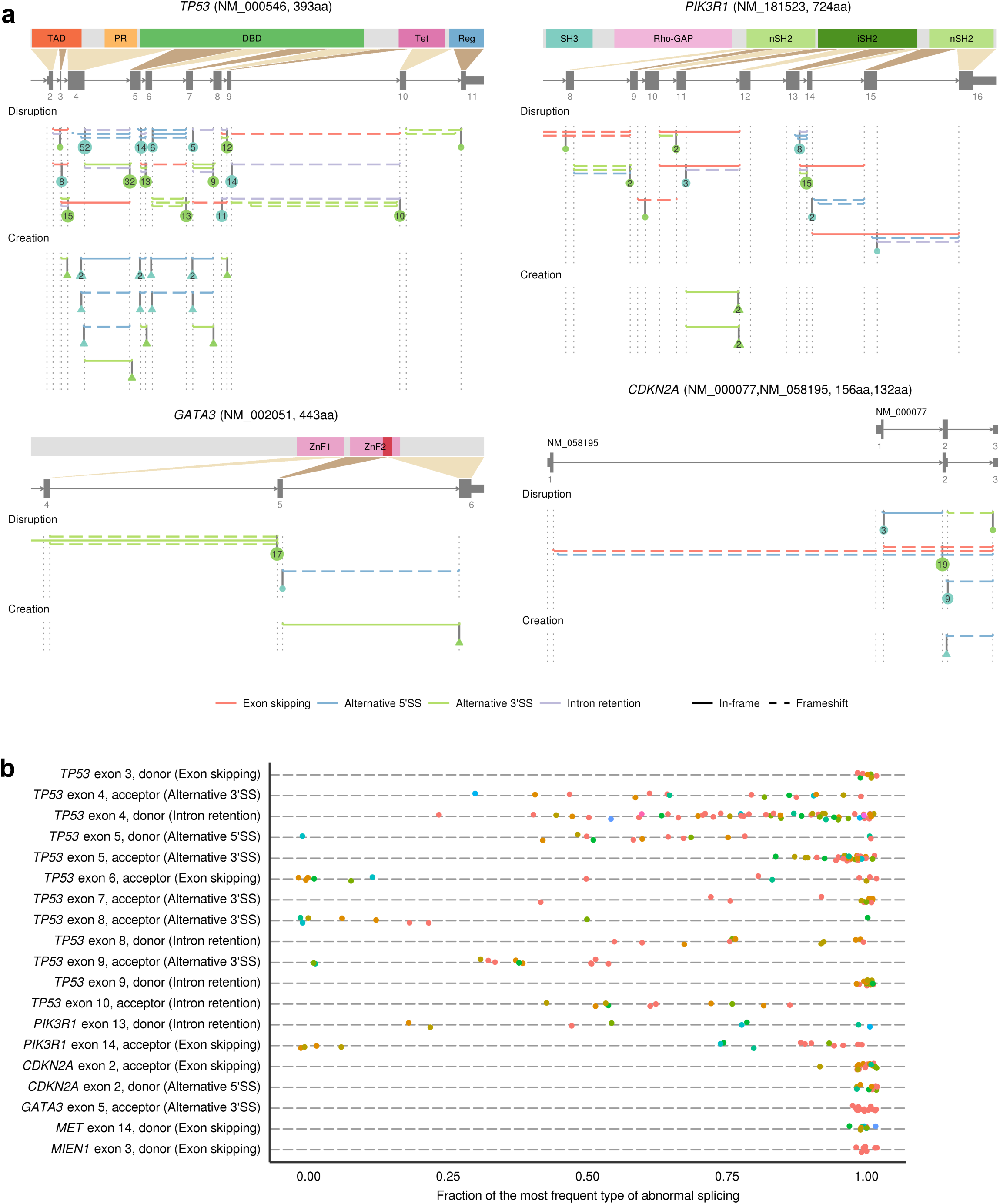
Genes frequently affected by SAVs in human cancers. **(a)** Distribution of SAVs and their resultant splicing outcomes for *TP53* (upper left), *PIK3R1* (upper right), *GATA3* (lower left), and *CDKN2A* (lower right). SS-disrupting and SS-creating SAVs are aggregated according to the authentic and alternative SSs, respectively. The number in circles or triangles represents the number of SS-disrupting and SS-creating SAVs for each SS, respectively. See also Supplementary Fig. 7a. **(b)** Fraction of the most frequent relative to total associated splicing outcomes for each SS-level SAV hotspot (found in ≥8 samples). The most frequent splicing outcome is noted in parentheses for each SS. The same color indicates the identical SAVs in terms of position and substitution or indel patterns. See also Supplementary Fig. 7b.

In many genes, particularly *NF1* and *RB1*, most SAVs and consequent splicing alteration events were diverse and widely distributed throughout the entire gene (Supplementary Fig. 7a, upper and middle), whereas several genes displayed prominent hotspots of SAVs (Supplementary Tables 6 and 7). Among the latter genes, SAVs affecting the same SSs tended to generate identical splicing consequences (Fig. 6b and Supplementary Fig. 7b). A typical example was *CDKN2A*, a well-known TSG that encodes the p16^INK4A^ and p14^ARF^ proteins, which was recurrently affected by SAVs targeting exon 2 common to both proteins. Other instances included *GATA3* SAVs found in breast invasive carcinomas (BRCA), in which most of them were the identical CA dinucleotide deletion at the acceptor site of exon 5, thus activating a cryptic 3’SS (7 nucleotides downstream), (Fig. 6a, lower left). Utilization of this cryptic splice acceptor caused a reading frameshift, resulting in loss of the second zinc finger (ZnF2) domain^28^. As was the case with *GATA3*, several genes showed tissue specificity, such as *FUBP1* and *ATRX* in lower grade gliomas (LGG), although most of the frequently altered genes were relevant across many cancer types (Fig. 5a).

Together with these well-established TSGs, SAVNet identified several recurrently altered genes (found in ≥10 samples) which had not been included in the cancer-related gene list^25^ but reported or predicted to function in a tumor suppressive manner, including *KANSL1*^29^, *NIPBL*^30^, *CUL3*^31^, *MYH9*^32^, *SMCHD1*^33^, and *HUWE1*^34^ (Fig. 5a). Thus, SAVNet may have potential to identify putative TSGs that are more prone to be affected by splicing aberrations. Conversely, *MET*, which encodes a hepatocyte growth factor receptor, was the only frequently affected oncogene, whose variants in the exon 14 donor site caused in-frame exon skipping known to activate c-Met^35^ (Fig. 5a and Supplementary Fig. 7a, lower left). Additionally, SAV hotspot analysis also identified recurrent SAVs occurring at the donor site of exon 3 of *MIEN1*, a putative oncogene located on the *HER2* amplicon^36^ (Fig. 6b and Supplementary Fig. 7a, lower right). Although the underlying mechanisms need to be clarified, SAVs may contribute to the activation of several oncogenes.

## Discussion

The development and application of SAVNet have led to the systematic detection of a substantial number of SAVs that had been overlooked by earlier studies^10,11^, although we focused only on those disrupting or creating splicing donor or acceptor motifs. Following previous studies^10,11,37^, our comprehensive and thorough analysis revealed the landscape of cis-acting somatic variants affecting splicing, characterized their positional differences and genomic features as well as underlying mutational processes in details, and argued that they include driver mutations, especially those inactivating TSGs. In particular, we demonstrated that exon skipping and alternative SS usage were more frequently involved in SAV-mediated TSG inactivation than intron retention. In addition, we clarified the relevance of SAVs at non-canonical sites, including the previously unrecognized position +3 and +5 of donor sites. The proposed framework with FDR control, which can dissect complex variant-splicing associations based on the Bayesian approach, is applicable to identify additional classes of somatic variants that disrupt splicing regulatory elements, including exonic and/or intronic splicing enhancers and silencers, although further elaboration of association rules will be required. Based on our findings, not only exonic but also intronic SNVs near exon-intron boundaries should be carefully evaluated as pathogenic variants, irrespective of the presence of amino acid changes. In the era of precision medicine, our framework will be critical to capture all driver variants, including previously overlooked SAVs, in cancer patients.

## URLs

SAVNet, https://github.com/friend1ws/SAVNet

Cancer Gene Census database, http://cancer.sanger.ac.uk/census

## Acknowledgments

This work was supported by Grant-in-Aid from the Japan Agency for Medical Research and Development [Advanced Genome Research and Bioinformatics Study to Facilitate Medical Innovation (17km0405207h0002)], Grant-in-Aid for Scientific Research (KAKENHI 15H05912, 15H05909, and 15K00398), and Post-K Research and Development (R&D) projects (hp170227). We thank Naoyuki Kataoka for expert opinion and Miki Sagou for technical assistance. The supercomputing resources were provided by the Human Genome Center, the Institute of Medical Science, The University of Tokyo. The results shown here are partly based upon data generated by the TCGA Research Network (http://cancergenome.nih.gov/).

## Author Contributions

Y.S. and K.K. designed the study. Y.S., K.C., A.O., H.T. and S.M. developed bioinformatics pipelines. Y.S. designed and implemented SAVNet. Y.S., K.K. and Y.K. analyzed and interpreted data. K.K. performed functional assays. Y.S. and K.K. generated figures and tables, and wrote the manuscript. S.O. and S.M. supervised the entire project. All authors participated in discussion and interpretation of the data and results.

## Methods

### Download of TCGA WES and RNA-seq data

WES and RNA-seq data were downloaded from Cancer Genomic Hub (https://cghub.ucsc.edu, currently hosted at NCI Genomic Data Commons (https://gdc-portal.nci.nih.gov/legacy-archive)). We used samples whose tumor and matched control WES and RNA-seq data are all available. We excluded LAML (acute myeloid leukemia) and OV (ovarian serous cystadenocarcinoma), because most of their DNA samples underwent whole genome amplification, leading to a large amount of artefactual variants.

### Alignment of TCGA WES data

As a reference genome, we used the sequences of assembled chromosomes, un-localized and un-placed scaffolds from GRCh37 (human reference assembly) as well as NC_007605 (Epstein-Barr virus) and hs37d5 (decoy from the 1000 Genomes Project Phase II) sequences. In WES analysis, for downloaded sequence data in BAM format, we first convert it to FASTQ format using bamtofastq command (with collate=1 exclude=QCFAIL,SECONDARY,SUPPLEMENTARY options) of biobambam (https://github.com/gt1/biobambam). FASTQ-formatted sequence were aligned with BWA-MEM version 0.7.8^38^ with −T0 option and sorted by biobambam bamsort command (with index=1 level=1 inputthreads=2 outputthreads=2 calmdnm=1 calmdnmrecompindentonly=1 options). Then, PCR duplicates were removed by biobambam bammarkduplicates command (with markthreads=2 rewritebam=1 rewritebamlevel=1 index=1 options).

### Detection of somatic SNVs and short indels

Our approach for detecting somatic SNVs and short indels consists of the following five steps:

1. Identification of candidate somatic SNVs and short indels using the approach based on Fisher’s exact test (as previously described^8^), which is currently implemented in GenomonFisher (https://github.com/Genomon-Project/GenomonFisher).
2. Excluding candidates present in pooled control samples by using EBFilter (https://github.com/Genomon-Project/EBFilter), a variant filtering algorithm based on a rigorous empirical Bayesian framework^39^.
3. Local re-alignment of short reads around candidate variants, which is implemented in GenomonMutationFilter (https://github.com/Genomon-Project/GenomonMutationFilter).
4. Removal of putative OxoG artifacts.
5. Annotation of the variants using Annovar^40^.

For step (1), we tallied up the numbers of mismatched bases at each position using short reads with mapping quality of ≥ 20 (for SNVs and indels) and with base quality of ≥ 15 (for SNVs). First, we roughly extracted the candidates satisfying the following criteria: (A) sequence depth ≥ 10, (B) mismatch ratio in tumor samples ≥ 0.05, (C) number of variant-supporting reads ≥ 4, and (D) mismatch ratio in matched control samples < 0.03. Next, we performed Fisher’s exact test to assess the differences in the ratios of the numbers of reference-supporting to variant-supporting reads between tumor and matched control samples, and candidate variants with P-value ≤ 0.1 were adopted.

For step (2), we performed filtering of all the remaining candidates, based on a beta-binomial error model, as described previously^39^. Briefly, we estimated the parameters of the beta-binomial error model using non-matched control samples (20 samples in this paper), obtained the predictive distributions of the mismatch ratios, and compared them with the observed mismatch ratio of tumor samples to quantify the statistical significance. We adopted candidate variants with P-value < 10^−4^.

For step (3), we performed local re-alignment of all short reads surrounding the candidate variants and their paired reads to the reference and variant-containing sequences, and counted the numbers of reference- and variant-supporting read pairs for tumor and matched control samples. We used “read pair-based” count to avoid double counting of a variant located in both reads of a single read pair with a small insert size. Then, we adopted candidates satisfying the following criteria: (A) number of variant-supporting read pairs in tumor samples ≥ 4, (B) number of variant-supporting read pairs in matched control samples ≤ 1, and (C) P-value of Fisher’s exact test comparing the ratios of the numbers of reference- and variant-supporting read pairs between tumor and matched control samples ≤ 0.1.

For step (4), to remove putative OxoG artifacts^41^, we calculated ALT_F1R2 (the number of variant-supporting read pairs whose first and second parts are aligned in the forward and reverse directions, respectively) and ALT_F2R1 (the number of variant-supporting read pairs whose first and second parts are aligned in the reverse and forward directions, respectively) for C > A and G > T substitutions. Then, C > A substitutions were removed if ALT_F1R2 < 2 or ALT_F2R1 / (ALT_F1R2 + ALT_F2R1) > 0.9, and G > T substitutions were removed if ALT_F2R1 < 2 or ALT_F1R2 / (ALT_F1R2 + ALT_F2R1) > 0.9.

### Alignment of TCGA RNA-seq data

Genome indexes were generated using STAR version 2.5.2a^42^ with the GRCh 37 release 19 GTF file (ftp://ftp.sanger.ac.uk/pub/gencode/Gencode_human/release_19/gencode.v19.annotation.gtf.gz) and –sjdbOverhang 100 option. For each sample, alignment to the reference genomes was performed by STAR version 2.5.2a with the following options: –runThreadN 6 –outSAMtype BAM Unsorted –outSAMstrandField intronMotif –outSAMunmapped Within –outSJfilterCountUniqueMin 1 1 1 1 –outSJfilterCountTotalMin 1 1 1 1 –outSJfilterOverhangMin 12 12 12 12 –outSJfilterDistToOtherSJmin 0 0 0 0 –alignIntronMax 500000 –alignMatesGapMax 500000 –alignSJstitchMismatchNmax −1 −1 −1 −1 –chimSegmentMin 12 –chimJunctionOverhangMin 12. Then, sorting and indexing of BAM files were performed using SAMtools version 1.2.

### Quantification of expression values for each gene from RNA-seq data

To quantify gene expression, we used our in-house software GenomonExpression (https://github.com/Genomon-Project/GenomonExpression), which calculates a slightly modified version of FPKM (fragments per kilobase of transcript per million mapped reads) measures^43^. Briefly, after excluding improperly aligned or low-quality read pairs (mapping quality < 20), sequence depth in the exonic regions was calculated, and normalized as per kilobase of exon as well as per million of aligned bases for each RefSeq gene. For genes with multiple transcript variants, their expression values were determined by selecting a transcript variant with the maximum FPKM value.

### Identification of splicing-associated variants (SAVNet)

To identify splicing-associated variants (SAVs), we developed and applied the novel approach, SAVNet (https://github.com/friend1ws/SAVNet), which consists of the following steps:

1. Collection of evidences of different types of abnormal splicing

We consider four types of abnormal splicing: exon skipping, alternative 5′ splice site (SS), alternative 3′SS usage and intron retention. The first three types (exon skipping, alternative 5′SS, alternative 3′SS) are extracted using splicing junctions (defined as pairs of start and end positions demarcated by spliced-aligned reads). We first extract abnormal splicing junctions (not registered in RefSeq genes (http://hgdownload.cse.ucsc.edu/goldenPath/hg19/database/refGene.txt.gz)) with ≥ 2 supporting reads (number of uniquely mapped reads crossing the junction) in at least one sample in the cohort by processing SJ.out.tab files generated as by-products of the STAR alignment step. Then, using our in-house program (junc_utils, https://github.com/friend1ws/junc_utils), we classify each splicing junction into exon skipping, alternative 5′SS or alternative 3′SS by the following criteria.

- Exon skipping: Two ends of the splicing junction correspond to annotated intron start (splicing donor) and end (splicing acceptor) sites of a gene, respectively.
- Alternative 5′SS: One end of the splicing junction corresponds to an annotated intron end (splicing acceptor) site of a gene, whereas the other end is located within the gene, but not at an annotated intron start (splicing donor) site of the gene.
- Alternative 3′SS: One end of the splicing junction corresponds to an annotated intron start (splicing donor) site of a gene, whereas the other end is located within the gene, but not at an annotated intron end (splicing acceptor) site of the gene.

Splicing junctions that do not meet any of the above are removed. Intron retentions are identified by our in-house program (intron_retention_utils simple_count command, https://github.com/friend1ws/intron_retention_utils). For each exon-intron boundary, the number of putative intron retention reads (those covering ≥ 10 bp of both sides of the exon-intron boundary) as well as that of normally spliced reads covering the last exonic base of the exon-intron boundary is counted. In this paper, to remove events observed in non-cancer samples, we used a panel of 742 control samples (collected from the TCGA cohort) and filtered out splicing junctions with ≥ 2 supporting reads in ≥ 8 control samples, and intron retentions whose intron retention fraction (the number of intron retention reads divided by total reads covering the exon-intron boundary) is ≥ 0.05 in ≥ 8 control samples.

2. Association of splicing alterations with somatic variants to construct possible variant-splicing bipartite graphs

In this step, we list up candidate combinations of somatic variants and possibly associated splicing alterations for each gene, which are subject to further investigation in the later step. For each gene, let ***g*** = (*g*_1_, *g*_2_,…,*gN*) *∊* {0,1,…,*M*}^*N*^ denote the status of somatic variants of *N* samples in the cohort, where *M* denotes the number of distinct somatic variants, *gN*= 0 represents that the *n*-th sample does not have any somatic variants in the target gene and *g*_*N*_ = *m* represents that *n* -th sample has the *m* -th somatic variant. Also, let 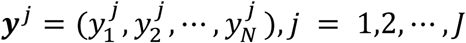, *j* = 1,2,…,*J* denote the number of supporting reads (the number of putative intron retention reads for intron retention) for the *j*-th splicing alteration, and let 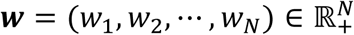 denote the weight for each sample used for normalization to negate variations in the amount of total sequence reads. We set *w*_*n*_ = *U*_*n*_/10^7^, where *U*_*n*_ is the number of uniquely aligned read pairs of the *n*-th sample.

The *m*-th somatic variant is considered to be associated with the *j*-th splicing alteration if the following three conditions are satisfied:

i. Their positional relationship implicates that the abnormal splicing can be a consequence of disruption of authentic SSs (those registered in the Refseq database) or creation of novel SSs caused by the somatic variant. More specifically, we check the following relationship:

- Abnormal splicing junction events caused by authentic SS disruption: (A) A somatic variant occurs at authentic splicing donor (between positions −3 (the third exonic base) through +6 (the sixth intronic base)) or acceptor sites (between −1 through +6), and (B) An abnormal splicing junction event (exon skipping and alternative 5′SS and 3′SS) encompass or are located 100 bp within the variant.
- Abnormal intron retention caused by authentic SS disruption: (A) A somatic variant occurs at authentic splicing donor or acceptor sites, and (B) an intron retention occurs at the disrupted SS or its opposite site of the same intron.
- Alternative SS usage caused by new SS creation: A somatic variant occurs within the newly created SS of an un-annotated junction end of an abnormal splicing event (alternative 5′SS or 3′SS).
ii. The average number of supporting reads for the *j*-th splicing alteration in samples with the *m*-th somatic variant is at least three times larger than those in samples without any somatic variants of the gene in consideration:

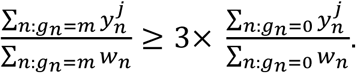
iii. The median number of supporting reads for the *j*-th splicing alteration in samples without any somatic variants of the gene in consideration is zero.

We create a bipartite graph (*V*_*M*_, *V*_*S*_, *E*) for the entire structure of variant-splicing associations, where vertices (*V*_*M*_, *V*_*S*_) represent somatic variants and splicing alterations, and edges (*E*) represent combinations of associated somatic variants and splicing alterations.

3. Pruning of edges to select the best model explaining the data

Here, we choose a sub-graph of the bipartite graph constructed in the previous step, which the most effectively explain the status of somatic variants and their impacts on splicing alterations (quantified by the numbers of supporting reads). We use the idea of “configuration” from previous eQTL and GWAS studies performed in complicated situations^12,13^. The configuration here is a |*E*|-dimensional binary vector ***γ*** = (*γ*_*m,j*_)_(*m,j*) *∊E*_, where *γ*_*m,j*_ *∊* {0,1} indicates whether the *m*-th variant and the *j***-**th splicing alterations have a causal relationship (1) or not (0). When there is no causal relationship between any somatic variants and splicing alterations (which we call the null model henceforth), ***γ*** = ***γ***^0^ where ***γ***^0^ is a vector whose elements are all zero 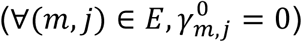. Under a configuration ***γ***, we classify somatic variants into “active” 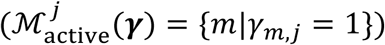 and “inactive” 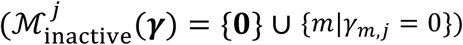 for the *j*-th splicing junction.

For each configuration ***γ***, we assume that the supporting reads ***y***^*j*^ are generated by Poisson distributions whose parameters are dependent on the activity status of somatic variants and multiplied by sample weights. The parameter of the Poisson distribution for the *n*-th sample is set to *w* _*N*_*λ*_0_ when it has only inactive variants on the *j*-th splicing alteration 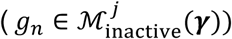, whereas it is set to *w*_*n*_*λ*_1_ for active variants 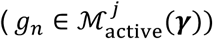 Additionally, we assume that *λ*_0_ and *λ*_1_ are generated by Gamma distribution with shape and rate parameters (*α*_0_, *β*_0_), (*α*_1_, *β*_1_), respectively. In this study, we set (*α*0, *β*_0_) = (1,1) and (*α*_1_, *β1*) = (1,0.01). Therefore, the likelihood of ***y***^*j*^ given ***γ*** is

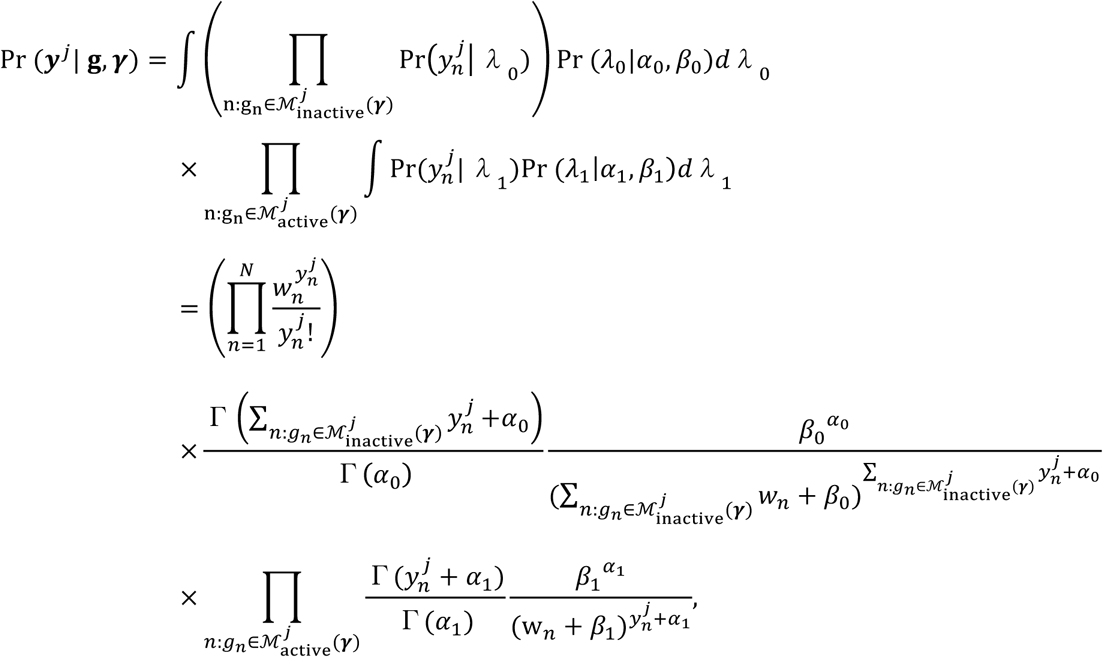

and the likelihood of whole the data (**Y** = {***y***^*j*^}_*j* = 1,2…,*j*_) 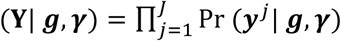. Also, the likelihood of ***y***^*j*^ under the null model ***γ*** = ***γ*^0^**, which can be calculated as a special case of the above, is

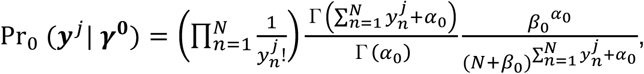

and the likelihood of whole the data is Pr_0_ 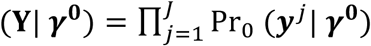.

For each variant *m*, we perform Bayesian model comparison to determine whether the somatic variant has any causal relationships with any splicing alterations (∃*j*, *γ*_*m,j*_ = 1) or not (∀*j*, *γ*_*m,j*_ = 0). Typically, to perform model comparison, we evaluate Bayes factor between the two distinct models. Here, as there are often many distinct null and non-null models, we aggregate these models through Bayesian Model Averaging and evaluate the Bayes factor between the aggregated null and non-null models:

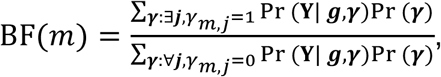

where Pr (***γ***) is set to be the uniform distribution. The variant *m* is identified as a SAV if its logarithm of Bayes factor is above the threshold (the default value is set to 3). Also, the splicing alterations caused by the variant *m* are identified by selecting the best model by 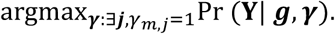.

4. Evaluation of FDR by permutation

To evaluate FDR, we permute the pairs of genomic and transcriptome data so that somatic variants and splicing alterations from different patients are coupled, and perform the same procedures (step 1 to 3). Assuming that *D*^target^ and 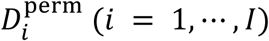 are the numbers of SAVs identified in the original step (correct combinations) and in the *i* -th permutation procedure, respectively, then FDR is estimated as

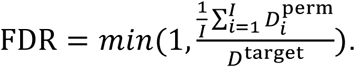

In this paper, we performed 100 permutation trials (*I* = 100).

5. Post-processing and rescuing SAVs

SAVs causing alternative intronic 5′ or 3′SSs are generally accompanied with intron retention at the original authentic SSs. Therefore, in these cases, we removed intron retention and retained only alternative 5′SS or 3′SSs in this paper. To sensitively detect recurrent SAVs, we performed additional screening and adopted variants satisfying the following criteria: (A) the combination of the same somatic variants (the same substitution at the same position) and the same splicing alterations was identified in other samples by the SAVNet procedure described above, (B) variant mismatch ratio in tumor samples ≥ 0.05, (C) number of variant-supporting reads ≥ 3, (D) mismatch ratio in tumor samples ≥ 10-fold of that in matched control samples, (E) number of reads supporting associated splicing alterations ≥ 2.

### Evaluation of influences of spliceosome variants on abnormal splicing

First, in the TCGA cohort, we searched for previously known somatic variants of splicing factors, including missense variants at K700, K666, H662, R625, E622, G740, G742, N626, and E902 of SF3B1, S34 and Q157 of U2AF1, and P95 of SRSF2, as well as missense, nonsense and frameshift variants of ZRSR2^7^. First, for each cancer type, we extracted splicing alterations with ≥ 2 supporting reads in at least one sample. Then, we identified splicing factors affected in ≥1% samples within each cancer type, and compared the number of RNA-seq reads supporting each splicing alteration between samples with and without the splice factor variants to derive P-value using t-test. Finally, we calculated Q-value for each splicing alteration using **qvalue** R package and splicing alterations with Q-value < 0.05 were considered to be associated with splice factor variants.

### Estimation of mutational signatures and membership of SAVs

We used **pmsignature** for estimating the signatures of mutational processes operative in the entire cohort and each cancer cohort as described in the previous paper^17^. Then, the extracted mutation signatures were classified into any of COSMIC signatures (http://cancer.sanger.ac.uk/cosmic/signatures) using minimum centered cosine similarity. Mutation signatures with centered cosine similarities to all the COSMIC signatures < 0.75 were classified to “other.” The estimates of membership (conditional probabilities attributed to each mutation signature) for each variant are provided by the following equation:

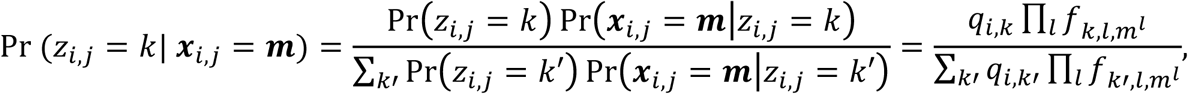

where the notation for each variable is described in the previous paper^17^. Finally, variant-level membership estimates were aggregated according to the COSMIC signatures and the presence of association with splicing alterations, so that the total numbers of variants and SAVs caused by each mutation signature (e.g., tobacco, ultraviolet) were estimated.

### Quantification of splicing-related features

MaxEnt^19^ and H-bond^20^ scores were calculated using **spliceSites** R package. To derive lengths and GC contents of exons affected by SAVs and their flanking introns, we extracted exonic nucleotides and adjacent upstream and downstream 150 intronic nucleotides. Then, we discarded 10 exonic and 20 intronic nucleotides from the exon-intron boundaries since they constitute splicing signals with specific nucleotides. Here, we excluded SS-disrupting SAVs affecting short exons (≤ 30bp), exons with multiple annotated start and end positions to avoid ambiguity.

### Cell line

HEK293T cells were obtained from RIKEN Cell Bank. Cell lines were authenticated by the provider and routinely tested for mycoplasma infection.

### Minigene splicing assay

For each region on interest, exonic and ∼300 bp flanking fragments containing either wild-type or variant sequences were synthesized (GeneArt, ThermoFisher Scientific) and cloned into the NheI and BamHI sites of the plasmid H492 (a kind gift from Prof. Masafumi Matsuo, Kobe University) using In-Fusion HD cloning kit (TaKaRa). Each construct was transiently transfected into HEK293T cells using X-tremeGENE 9 DNA Transfection Reagent (Roche) in six-well tissue culture plates. Forty-eight hours after transfection, total RNA was isolated with RNeasy Mini kit (QIAGEN), and used to synthesize cDNA with ReverTra Ace qPCR RT Kit (TOYOBO). Each cDNA was amplified with KOD FX Neo DNA polymerase (TOYOBO) using the primers (forward, 5′-ATTACTCGCTCAGAAGCTGTGTTGC-3′, and reverse, 5′-AAGTCTCTCACTTAGCAACTGGCAG-3′), which correspond to sequences of exons of H492. PCR products were separated by electrophoresis on 2% agarose gel and visualized with a UV transilluminator (UVP). To confirm the sequence of each band, the PCR products were gel purified and analyzed by Sanger sequencing.

### Data analysis

All analyses were performed in Python 2.7.8 and R 3.3.2 (R Core Team) and most figures were generated using the **ggplot2** R package. In all box plots, the center line and lower and upper hinges correspond to the median, and the first and third quartiles (25 and 75 percentiles), respectively. The upper and lower whiskers extend from the upper and lower hinges to the largest or smallest values no further than 1.5 * IQR from the hinges, respectively, where IQR represents inter-quartile range, or distance between the first and third quartiles. Sequence logos were drawn via our in-house program (https://github.com/friend1ws/ggseqlogo).

### Data availability

The raw sequence data used in this study can be downloaded by registered users from https://gdc-portal.nci.nih.gov/. The processed data and scripts for generating figures are available from the corresponding author upon reasonable request.

## Supplementary figure legends

**Supplementary Figure 1: Evaluation of the performance of SAVNet.**

**(a)** An illustration of a simulation study on the effect of the number of potentially associated splicing alterations on the performance of SAV detection. The numbers of splicing-supporting reads are generated by Poisson distribution with parameters of 3 or 0.3 when splicing alterations are actually associated (true) or not associated (false) with a somatic variant, respectively. Here, we assume an all-or-nothing situation: either all possible splicing alterations or none are associated with a somatic variant. For each group, 100 trials with 20 samples were performed. **(b)** Box plots showing the logarithm of the Bayes factor for each number of true (red) and false (gray) variant-splicing associations. Also, the sensitivity (the fraction of trials where the logarithm of the Bayes factor was greater than 3.0 and the default threshold used in this study) is shown in purple points with lines for the true association group. As the number of associated splicing alterations increases, the sensitivity improves dramatically. On the other hand, the logarithm of the Bayes factor for the false association group does not reach the threshold irrespective of the number of associated splicing alterations, indicating the high specificity of SAVNet. **(c)** FDR for each cancer type estimated by permutation of combinations of WES and RNA-seq data when the threshold of the logarithm of the Bayes factor was set to 3.0. For ESCA and STAD, the thresholds were adjusted so that their FDRs were maintained below 0.05. **(d, e)** Venn diagrams showing the overlap of SAVs detected by the current (SAVNet) and previous (ratio-based splicing analysis) studies^10^. SAVs at positions analyzed by both studies (−1, +1, and +2 of donor and acceptor sites) were evaluated for all genes **(d)** and candidate cancer-related genes^25^ **(e)**.

**Supplementary Figure 2: Examples of SAVs associated with multiple (≥ 5) splicing alterations.**

Different colors indicate different types of splicing alterations. The solid and dashed lines represent in-frame and frameshift alterations, respectively. The number at each line represents the count of supporting reads.

**Supplementary Figure 3: Positional distribution and underlying mutational processes of SAVs.**

**(a)** Number of splicing-associated indels disrupting splice donor and acceptor sites (upper) and their fraction relative to total indels (lower) at canonical or non-canonical sites. **(b)** Visualization of mutational patterns for five representative mutation signatures identified by pmsignature software^17^. These patterns are defined by substitution type, strand bias, and sequence context of 2 bp within the substituted bases, as described in ref. 17. **(c)** Fraction of estimated SAVs relative to estimated total variants attributed to each mutational signature in each cancer type. Red dashed lines represent the overall fraction of SAVs relative to total variants for each cancer. **(d)** Two typical examples of SS-creating SAVs (through formations of GT and YAG motifs, respectively) are displayed. **(e)** The effects of SAVs on splicing strength (based on MaxEnt or H-bond scores) for authentic or alternative SSs were assessed. For SS-disrupting SAVs, the difference in splicing strength between alternative and unsubstituted authentic SSs (WT) was compared with that between alternative and substituted authentic SSs (Mut). For SS-creating SAVs, the difference between unsubstituted alternative and authentic SSs (WT) was compared with that between substituted alternative and authentic SSs (Mut). **(f, g)** Box plots showing the differences of MaxEnt scores for alternative 5′SSs and 3′SSs **(f)** and H-bond scores for alternative 5′SSs **(g)**.

**Supplementary Figure 4: Genomic features of SSs affected by SS-disrupting SAVs based on their splicing outcomes.**

**(a)** Schematics depicting the classification methods for somatic variants affecting authentic SSs by their splicing outcomes**. (b, c)** Changes in MaxEnt scores by somatic variants were compared between normal and abnormal splicing groups according to the substituted position of donor **(b)** or acceptor **(c)** sites. “Normal splicing” represents samples lacking the relevant splicing alterations in the presence of somatic variants. **(d)** Change in H-bond scores triggered by somatic variants at authentic splicing donor sites according to splicing outcomes. **(e)** Changes in H-bond scores by somatic variants were compared between normal and abnormal splicing groups according to the substituted position of donor sites. **(f)** Sequence motifs of splicing acceptor sites at which somatic SNVs lead to normal (left) or abnormal splicing (right; identified by SAVNet) according to the variant position. **(g)** Sequence motifs of splicing donor sites at which somatic SNVs lead to normal splicing, exon skipping, alternative 5′SS, intron retention, or complex abnormalities. Categories whose number of SAVs is less than 25 are not displayed.

**Supplementary Figure 5: The exon-intron architecture affected by SS-disrupting SAVs based on their splicing outcomes.**

**(a)** Lengths of exons affected by SAVs, and their 5′ and 3′ introns were compared among the five splicing groups. **(b)** MaxEnt scores (before substitution) at authentic splicing donor (left) and acceptor (right) sites according to splicing outcomes. **(c)** H-bond scores (before substitution) at authentic splicing donor sites according to splicing outcomes. **(d-f)** Matrices of the significance of differences in GC content **(d)**, length **(e)** of exons affected by SAVs and their 5′ and 3′ introns, and splicing strength (based on MaxEnt or H-bond scores) of affected SSs **(f)** among the splicing groups. The significant differences were quantified by the negative logarithm of the P-value of one-sided Wilcoxon rank-sum tests whose alternative hypotheses are that either the splicing pattern 1 is less (blue) or greater (red) than splicing pattern 2. The smaller P-value between the two alternative hypotheses is shown.

**Supplementary Figure 6: Enrichment of SAVs in cancer-related genes.**

**(a)** The fraction of SAVs affecting cancer-related genes relative to total SAVs according to splicing outcomes were compared with other types of somatic variants (silent, missense, and nonsense). These cancer-related gene sets are derived from three publications (refs. 1, 25, and 26) and the Cancer Gene Census database (CGC, as of Feb. 2017). **(b)** The fraction of somatic variants affecting oncogenes or TSGs (ref. 1) for each type of abnormal splicing was compared with that of silent variants. The bar plot shows the logarithm of P-values calculated using a binomial test.

**Supplementary Figure 7: Genes frequently affected by SAVs in human cancers.**

**(a)** Distribution of SAVs and their resultant splicing outcomes for *NF1* (upper), *RB1* (middle), *MET* (lower left), and *MIEN1* (lower right). SS-disrupting SAVs are aggregated according to the authentic SSs. The number in circles represents the number of SAVs for each SS. **(b)** Schematics depicting methods to quantify the fraction of the most frequent relative to total associated splicing outcomes for each SS-level SAV hotspot. The number at each line represents the count of supporting reads.

## References

1 Vogelstein, B. et al. Cancer genome landscapes. Science 339, 1546–1558, doi:10.1126/science.1235122 (2013).

2 Garraway, L. A. & Lander, E. S. Lessons from the cancer genome. Cell 153, 17–37, doi:10.1016/j.cell.2013.03.002 (2013).

3 Martincorena, I. & Campbell, P. J. Somatic mutation in cancer and normal cells. Science 349, 1483–1489, doi:10.1126/science.aab4082 (2015).

4 Kalnina, Z., Zayakin, P., Silina, K. & Line, A. Alterations of pre-mRNA splicing in cancer. Genes Chromosomes Cancer 42, 342–357, doi:10.1002/gcc.20156 (2005).

5 Venables, J. P. Aberrant and alternative splicing in cancer. Cancer Res 64, 7647–7654, doi:10.1158/0008-5472.CAN-04-1910 (2004).

6 Scotti, M. M. & Swanson, M. S. RNA mis-splicing in disease. Nat Rev Genet 17, 19–32, doi:10.1038/nrg.2015.3 (2016).

7 Dvinge, H., Kim, E., Abdel-Wahab, O. & Bradley, R. K. RNA splicing factors as oncoproteins and tumour suppressors. Nat Rev Cancer 16, 413–430, doi:10.1038/nrc.2016.51 (2016).

8 Yoshida, K. et al. Frequent pathway mutations of splicing machinery in myelodysplasia. Nature 478, 64–69, doi:10.1038/nature10496 (2011).

9 Brooks, A. N. et al. A pan-cancer analysis of transcriptome changes associated with somatic mutations in U2AF1 reveals commonly altered splicing events. PLoS One 9, e87361, doi:10.1371/journal.pone.0087361 (2014).

10 Jung, H. et al. Intron retention is a widespread mechanism of tumor-suppressor inactivation. Nat Genet 47, 1242–1248, doi:10.1038/ng.3414 (2015).

11 Supek, F., Miñana, B., Valcárcel, J., Gabaldón, T. & Lehner, B. Synonymous mutations frequently act as driver mutations in human cancers. Cell 156, 1324–1335, doi:10.1016/j.cell.2014.01.051 (2014).

12 Stephens, M. A unified framework for association analysis with multiple related phenotypes. PLoS One 8, e65245, doi:10.1371/journal.pone.0065245 (2013).

13 Flutre, T., Wen, X., Pritchard, J. & Stephens, M. A statistical framework for joint eQTL analysis in multiple tissues. PLoS Genet 9, e1003486, doi:10.1371/journal.pgen.1003486 (2013).

14 Sibley, C. R., Blazquez, L. & Ule, J. Lessons from non-canonical splicing. Nat Rev Genet 17, 407–421, doi:10.1038/nrg.2016.46 (2016).

15 Lee, Y. & Rio, D. C. Mechanisms and Regulation of Alternative Pre-mRNA Splicing. Annu Rev Biochem 84, 291–323, doi:10.1146/annurev-biochem-060614-034316 (2015).

16 Nishida, A. et al. Chemical treatment enhances skipping of a mutated exon in the dystrophin gene. Nat Commun 2, 308, doi:10.1038/ncomms1306 (2011).

17 Shiraishi, Y., Tremmel, G., Miyano, S. & Stephens, M. A Simple Model-Based Approach to Inferring and Visualizing Cancer Mutation Signatures. PLoS Genet 11, e1005657, doi:10.1371/journal.pgen.1005657 (2015).

18 Alexandrov, L. B. et al. Signatures of mutational processes in human cancer. Nature 500, 415–421, doi:10.1038/nature12477 (2013).

19 Yeo, G. & Burge, C. B. Maximum entropy modeling of short sequence motifs with applications to RNA splicing signals. J Comput Biol 11, 377–394, doi:10.1089/1066527041410418 (2004).

20 Freund, M. et al. A novel approach to describe a U1 snRNA binding site. Nucleic Acids Res 31, 6963–6975 (2003).

21 Carmel, I. Comparative analysis detects dependencies among the 5’ splice-site positions. Rna 10, 828–840, doi:10.1261/rna.5196404 (2004).

22 Burge, C. & Karlin, S. Prediction of complete gene structures in human genomic DNA. J Mol Biol 268, 78–94, doi:10.1006/jmbi.1997.0951 (1997).

23 Naftelberg, S., Schor, I. E., Ast, G. & Kornblihtt, A. R. Regulation of alternative splicing through coupling with transcription and chromatin structure. Annu Rev Biochem 84, 165–198, doi:10.1146/annurev-biochem-060614-034242 (2015).

24 Keren, H., Lev-Maor, G. & Ast, G. Alternative splicing and evolution: diversification, exon definition and function. Nat Rev Genet 11, 345–355, doi:10.1038/nrg2776 (2010).

25 Ye, K. et al. Systematic discovery of complex insertions and deletions in human cancers. Nat Med 22, 97–104, doi:10.1038/nm.4002 (2016).

26 Lawrence, M. S. et al. Discovery and saturation analysis of cancer genes across 21 tumour types. Nature 505, 495–501, doi:10.1038/nature12912 (2014).

27 Cheung, L. W. et al. High frequency of PIK3R1 and PIK3R2 mutations in endometrial cancer elucidates a novel mechanism for regulation of PTEN protein stability. Cancer Discov 1, 170–185, doi:10.1158/2159-8290.CD-11-0039 (2011).

28 Usary, J. et al. Mutation of GATA3 in human breast tumors. Oncogene 23, 7669–7678, doi:10.1038/sj.onc.1207966 (2004).

29 Yoshida, K. et al. The landscape of somatic mutations in Down syndrome-related myeloid disorders. Nat Genet 45, 1293–1299, doi:10.1038/ng.2759 (2013).

30 Barber, T. D. et al. Chromatid cohesion defects may underlie chromosome instability in human colorectal cancers. Proc Natl Acad Sci U S A 105, 3443–3448, doi:10.1073/pnas.0712384105 (2008).

31 Ooi, A. et al. CUL3 and NRF2 mutations confer an NRF2 activation phenotype in a sporadic form of papillary renal cell carcinoma. Cancer Res 73, 2044–2051, doi:10.1158/0008-5472.CAN-12-3227 (2013).

32 Schramek, D. et al. Direct in vivo RNAi screen unveils myosin IIa as a tumor suppressor of squamous cell carcinomas. Science 343, 309–313, doi:10.1126/science.1248627 (2014).

33 Leong, H. S. et al. Epigenetic regulator Smchd1 functions as a tumor suppressor. Cancer Res 73, 1591–1599, doi:10.1158/0008-5472.CAN-12-3019 (2013).

34 Inoue, S. et al. Mule/Huwe1/Arf-BP1 suppresses Ras-driven tumorigenesis by preventing c-Myc/Miz1-mediated down-regulation of p21 and p15. Genes Dev 27, 1101–1114, doi:10.1101/gad.214577.113 (2013).

35 Ma, P. C. et al. Functional expression and mutations of c-Met and its therapeutic inhibition with SU11274 and small interfering RNA in non-small cell lung cancer. Cancer Res 65, 1479–1488, doi:10.1158/0008-5472.CAN-04-2650 (2005).

36 Dasgupta, S. et al. Novel gene C17orf37 in 17q12 amplicon promotes migration and invasion of prostate cancer cells. Oncogene 28, 2860–2872, doi:10.1038/onc.2009.145 (2009).

37 Xu, Q. & Lee, C. Discovery of novel splice forms and functional analysis of cancer-specific alternative splicing in human expressed sequences. Nucleic Acids Res 31, 5635–5643 (2003).

38 Li, H. & Durbin, R. Fast and accurate short read alignment with Burrows-Wheeler transform. Bioinformatics 25, 1754–1760, doi:10.1093/bioinformatics/btp324 (2009).

39 Shiraishi, Y. et al. An empirical Bayesian framework for somatic mutation detection from cancer genome sequencing data. Nucleic Acids Res 41, |pe89, doi:10.1093/nar/gkt126 (2013).

40 Wang, K., Li, M. & Hakonarson, H. ANNOVAR: functional annotation of genetic variants from high–throughput sequencing data. Nucleic Acids Res 38, e164, doi:10.1093/nar/gkq603 (2010).

41 Costello, M. et al. Discovery and characterization of artifactual mutations in deep coverage targeted capture sequencing data due to oxidative DNA damage during sample preparation. Nucleic Acids Res 41, e67, doi:10.1093/nar/gks1443 (2013).

42 Dobin, A. et al. STAR: ultrafast universal RNA–seq aligner. Bioinformatics 29, 15–21, doi:10.1093/bioinformatics/bts635 (2013).

43 Shiraishi, Y. et al. Integrated analysis of whole genome and transcriptome sequencing reveals diverse transcriptomic aberrations driven by somatic genomic changes in liver cancers. PLoS One 9, e114263, doi:10.1371/journal.pone.0114263 (2014).

